# vWCluster: A Network Based Clustering of Multi-omics Breast Cancer Data Based on Vector-Valued Optimal Transport

**DOI:** 10.1101/2021.06.17.448878

**Authors:** Jiening Zhu, Jung Hun Oh, Joseph O. Deasy, Allen Tannenbaum

**Affiliations:** Department of Applied Mathematics & Statistics, Stony Brook University, NY; Department of Medical Physics, Memorial Sloan Kettering Cancer Center, NY; Departments of Computer Science and Applied Mathematics & Statistics, Stony Brook University, NY

## Abstract

In this paper, we present a network-based clustering method based on the vector-valued Wasserstein distance derived from optimal mass transport (OMT) theory. This distance allows for the natural integration of multi-layer representations of data in a given network from which one derives clusters via a hierarchical clustering approach. In this study, we applied the methodology, called ***vector Wasserstein clustering*** (vW-cluster), to multi-omics data from the two largest breast cancer studies. The resultant clusters showed significantly different survival rates in Kaplan-Meier analysis in both datasets. CIBERSORT scores were compared among the identified clusters. Out of the 22 CIBERSORT immune cell types, 9 were commonly significantly different in both datasets, suggesting the difference of tumor immune microenvironment in the cluster. vWCluster can aggregate multi-omics data represented as a vectorial form in a network with multiple layers, taking into account the concordant effect of heterogeneous data, and further identify subtypes of tumors with different survival rates.

## 1 Introduction

Current large-scale cancer genome projects, such as The Cancer Genome Atlas (TCGA), provide comprehensive molecular portrait of human cancers, including gene expression, copy number variation (CNV), and DNA methylation profiles [1]. These offer unprecedented opportunities for exploring cancer biology that is characterized through various molecular functions and their complex interactions. Several clustering methods have been proposed to identify tumor subtypes associated with distinct clinical outcomes, leveraging complementary multi-omics data [16]. iCluster uses a joint latent variable method across multi-omics types to model integrative clustering [27]. Recently, network based methods on multi-omics data have been proposed. Similarity network function (SNF) is a technique for combining multiple networks, which of each indicates the similarity of samples based on each omics type, into a single network, subsequently followed by spectral clustering to identify sub-types of tumors [30]. On the other hand, aWCluster utilizes a prior known network of gene products to integrate multi-omics data [23]. First, the data integration method yields an invariant measure for each node (gene). The Wasserstein distance, derived from optimal mass transport (OMT) theory [28, 29], is then computed between all pairs of samples on the network using the invariant measure. The resulting distance matrix is then input to a hierarchical clustering, resulting in subtypes.

The 1-Wasserstein distance, also known as the Earth Mover’s Distance (EMD), was first formulated by the French civil engineer and mathematician Gaspard Monge in 1781 [13, 25, 28, 29]. Originally, this subject was inspired by the problem of finding the optimal plan, relative to a given cost, for moving a pile of soil from a given location to another in a mass preserving manner. The next major breakthrough is due to Leonid Kantorovich [17], who relaxed the original problem and introduced linear programming for its solution. As a natural tool to analyze data distributions, OMT is becoming more and more widely used in signal processing, machine learning, computer vision, meteorology, statistical physics, quantum mechanics, and network theory [3, 4, 15, 19, 20, 26]. To even strengthen its power, several works deal with extensions of the theory to the vector-valued and even matrix cases; see [4, 5, 8, 9, 10, 20] and the references therein.

In the current study, we propose a new method, called ***vector Wasserstein clustering*** (vWCluster), in which we employ a vector-valued version of the Wasserstein distance [11]. The method is applied to the two largest breast cancer datasets from the Molecular Taxonomy of Breast Cancer International Consortium (METABRIC) and TCGA studies [12, 18]. First, each data type in multi-omics data is represented as a layer of a biological network. The Wasserstein distance is then computed on the vector-valued data in the network between all pairs of samples. The resulting distance matrix is then inputted into a hierarchical clustering method to identify various subtypes of tumors. In the following section, we describe the proposed method and data in detail.

## 2 Methods

We developed a vector-valued OMT approach that integrates multi-omics data represented in a multi-layer network. The resulting distance matrix is used as the input to a clustering method to identify subgroups of samples. This method is based on the 𝒲_1_ Wasserstein distance (as known as EMD). Accordingly, we will only outline the OMT theory in this special case. In the following, we first describe the basic concept of Wasserstein distance and then introduce the proposed method.

### 2.1 Scalar-valued optimal transport

The original Monge’s formulation of OMT (in which the cost function is defined by the distance) [28, 29] may be given a modern expression as follows:

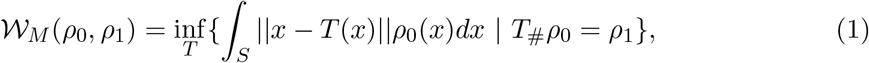

where *S* denotes a subdomain of ℝ^*n*^, *T* is the transport map, and *ρ*_0_, *ρ*_1_ are two probability distributions. Here *T*_#_ denotes the push-forward of *T*. The *Wasserstein 1-distance* is the optimal cost with respect to the norm among all possible *T*. Note that mass ia preserved in Monge’s formulation. Since we will only be using the distance as the cost for simplicity, by *Wasserstein distance*, we will always mean *Wasserstein 1-distance*. This is also called the Earth Mover’s distance (EMD).

As pioneered by Leonid Kantorovich [17], the Monge formulation of OMT may be relaxed by replacing transport maps *T* by couplings *π*:

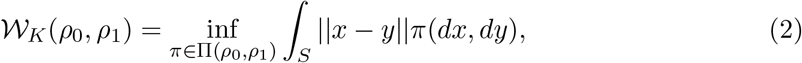

where Π(*ρ*_0_, *ρ*_1_) denotes the set of all the couplings between *ρ*_0_ and *ρ*_1_ (joint distributions whose two marginal distributions are *ρ*_0_ and *ρ*_1_). Despite of the relaxation, one may show Kantorovich and Monge formulations are equivalent in a number of cases under certain continuity constraints; see [28, 29] and the references therein.

One of the benefits of (2) is that it amounts to a linear programming problem whose dual is easy to compute. Dualizing again, an equivalent form may be expressed as follows (see [13] for the proof):

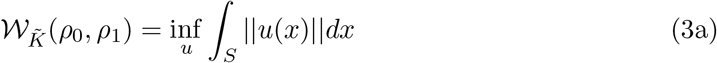

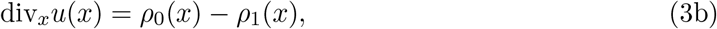

where *u* = (*u*_1_, *u*_2_, …, *u*_*n*_) : *S* →ℝ^*n*^ is the flux, and div_*x*_ denotes the divergence operator. It is straightforward to extend (3) to the discrete case by simply replacing the integral by an appropriate summation and replacing div_*x*_ by the discrete divergence operator:

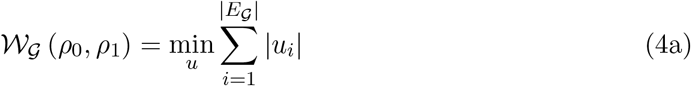

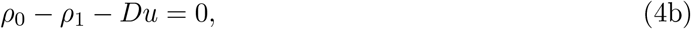

On the graph 𝒢 = (*V*_*𝒢*_, *E*_*𝒢*_), the fluxes *u*_*i*_ now are defined on the edges *E*_*𝒢*_, and 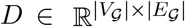 denotes the incidence matrix of 𝒢 with directionality, namely we need to specify the directions of the fluxes. Thus in the matrix *D* each column has two nonzero entries, where one is 1 whose row number is the starting point of an edge while the other nonzero entry is -1 whose row number is the ending point of that edge.

### 2.2 Vector-valued optimal transport

A vector-valued density 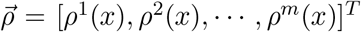 on a given space *S* (continuous or discrete) may represent a physical entity that can mutate or transition between alternative manifestations, e.g., power reflected off a surface at different frequencies or polarizations. More formally, an *m*-layer *vector-valued density* 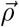 on *S* ⊆ℝ^*n*^ is a map from *S* to 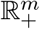 whose total mass is defined as 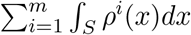. As a distribution, we require that the total mass should be 1.

Vector-valued optimal transport studies such distributions, which is of great theoretical and practical interest since it does not simply consider each layer separately, but explicitly models the the relationships among layers. A relationship is expressed as an additional graph structure that connects each layer. Specifically, each component of *ρ* is represented by a node of a graph ℱ = (*V*_*ℱ*_, *E*_*ℱ*_) and an edge between two nodes allows for direct transport between the corresponding layers. So |*V*_*ℱ*_ | = *m*, which represents the cardinality of all the channels, and *E*_*F*_ is the set of all the direct connections of between the layers.

Thus, the vector-valued optimal transport problem may be written as follows:

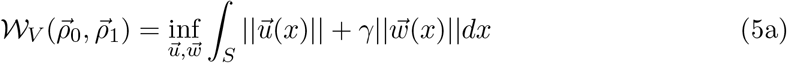

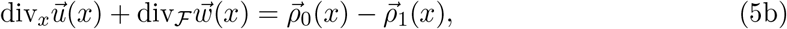

where 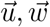 are both vector-valued, div_*x*_ is the spatial divergence which is taken component-wise for each layer, and div_*ℱ*_ is the discrete divergence on the graph ℱ which takes the flows between channels into account. Here *γ* ≥ 0 is parameter to control flow between channels.

As in the scalar-valued case, we can extend the definition for distributions to a discrete graph 𝒢. The vector-valued formulation on a graph is then the following:

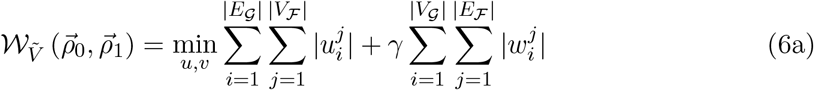

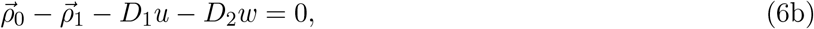

where *u* is the flux within each layer, *w* is the flux cross layers, and *D*_1_ and *D*_2_ are two matrices of the discrete divergence operators for two graphs.

On the one hand, this is a generalized form of (4) derived by replacing each original node in the graph 𝒢 by another graph. On the other hand, this formulation may be understood as a distribution on a super-graph 𝒢 ×ℱ. This super-graph is a irregular grid version of the Kronecker product. A slight difference from directly computing OMT distance on such a super-graph is that vector OMT on a graph here gives two different weights for the two different kinds of edges. It is weighted vector-valued OMT. We later will see, that two different kinds of fluxes via two graphs have different meanings.

### 2.3 Multi-omics data from two breast cancer studies

Multi-omics data for METABRIC and TCGA breast cancer studies were downloaded from the cBioPortal database [6, 14]. The METABRIC dataset contains microarray gene expression of 24,368 genes from 1,904 samples and copy number variation (CNV) of 22,544 genes from 2,173 samples. The intersection of the two omics data resulted in 16,195 genes from 1,904 samples. The TCGA breast cancer dataset consists of RAN-Seq gene expression of 18,022 genes from 1,100 samples, CNV of 15,213 genes from 1,080 samples, and methylation of 15,585 genes from 741 samples. The intersection of the three omics data resulted in 7,737 genes from 726 samples.

### 2.4 Graph structures for analysis

We represented multi-omics data as vector-valued distributions on the gene (product) interaction network. The interaction network was derived from HPRD[24]. The largest connected network component was found in the interaction of the HPRD and the gene list of METABRIC or TCGA breast cancer data, separately, resulting in 3147 genes and 3426 genes, respectively. As multi-omics data in the TCGA breast cancer cohort, gene expression, CNV, and methylation data were used, whereas in the METABRIC cohort, only gene expression and CNV data were available, thereby forming 3-vector and 2-vector distributions, respectively.

The network for METABRIC consisted of two layers (gene expression and CNV), each of which had the same topology (the largest connected network component) derived from HPRD. The connection between the two layers was formed by connecting the two nodes for the same gene in each layer, yielding the graph ℱ structure.

For the network with the TCGA data, the layer for gene expression was connected with both layers for CNV and methylation since CNV and methylation may affect the level of RNA gene expression. There was no connection between the CNV and methylation layers.

### 2.5 Markov chain and stationary distribution

One problem of applying the vector-valued optimal transport method to multi-omics data is that the scale of individual omics data varies. For example, CNV data consists of integer values, while gene expression and methylation data have continuous values. To tackle this issue, we use the invariant (stationary) distribution derived from a Markov process of the gene network.

A *Markov process* is a stochastic process such that the probability of a given event depends only on the state of the previous event To put it simpler in our graph setting, one starts with a certain distribution. At each time step, the probability at each node redistributes to all its neighbors with predefined weights. In the gene network setting [7], we set the probability of moving from a node *i* to its neighbor *j* to be:

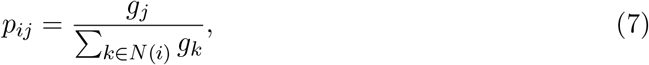

where *g*_*k*_ is the weight of node k, which can be either of the any omics type (gene expression, CNV or methylation). Note that for methylation, 1-methylation values were used since methylation is likely to be negatively correlated with gene expression.

The matrix *p* is a stochastic matrix, i.e., the state probability matrix from the current time step to the next as follows:

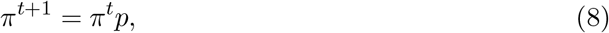

where *π*^*t*^ is the distribution at time step *t*. In our setting, after a finite number of time steps, the initial distribution will converge to a stationary (invariant) distribution *π* such that

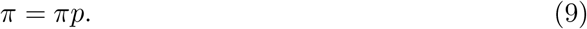

The stationary distribution has a closed form solution:

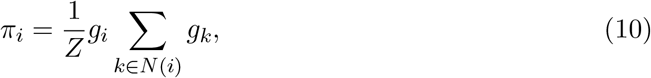

where *Z* is the normalization factor to be a probability distribution.

This Markov process on the gene network mimics the interactions among genes and the stationary distribution gives a distribution that represents the information each gene has which includes not only its own value but the interactions with its neighbors. The Markov process was performed for each sample in individual omics types, separately, yielding invariant measures *I*_*ijk*_, which are invariant measures for sample *i*, omics type *j*, and gene *k*.

### 2.6 Clustering based on the vector-valued Wasserstein distance

With the graph structure determined, the vector-valued Wasserstein distance was computed for each pair of samples, using the invariant measures of each omics type. Note that the network for METABRIC or TCGA breast cancer data consisted of 2 or 3 layers, respectively. That is, we fitted the multi-omics data into the vector-valued optimal transport model. The resulting distance matrix was then input to standard hierarchical clustering to identify clusters of tumors. Kaplan-Meier survival analysis with log-rank test was performed to assess the difference of survival rates among the clusters identified. Further, CIBERSORT scores were compared among the clusters [2, 21]. This analysis was performed for METABRIC or TCGA breast cancer data, separately.

## 3 Results

### 3.1 METABRIC data analysis

The vector-valued Wasserstein distance was computed on gene expression and CNV data for METABRIC data. As above, the resulting distance matrix was input to standard hierarchical clustering. The clustering results are shown in Figure 2.

**Figure 1:**
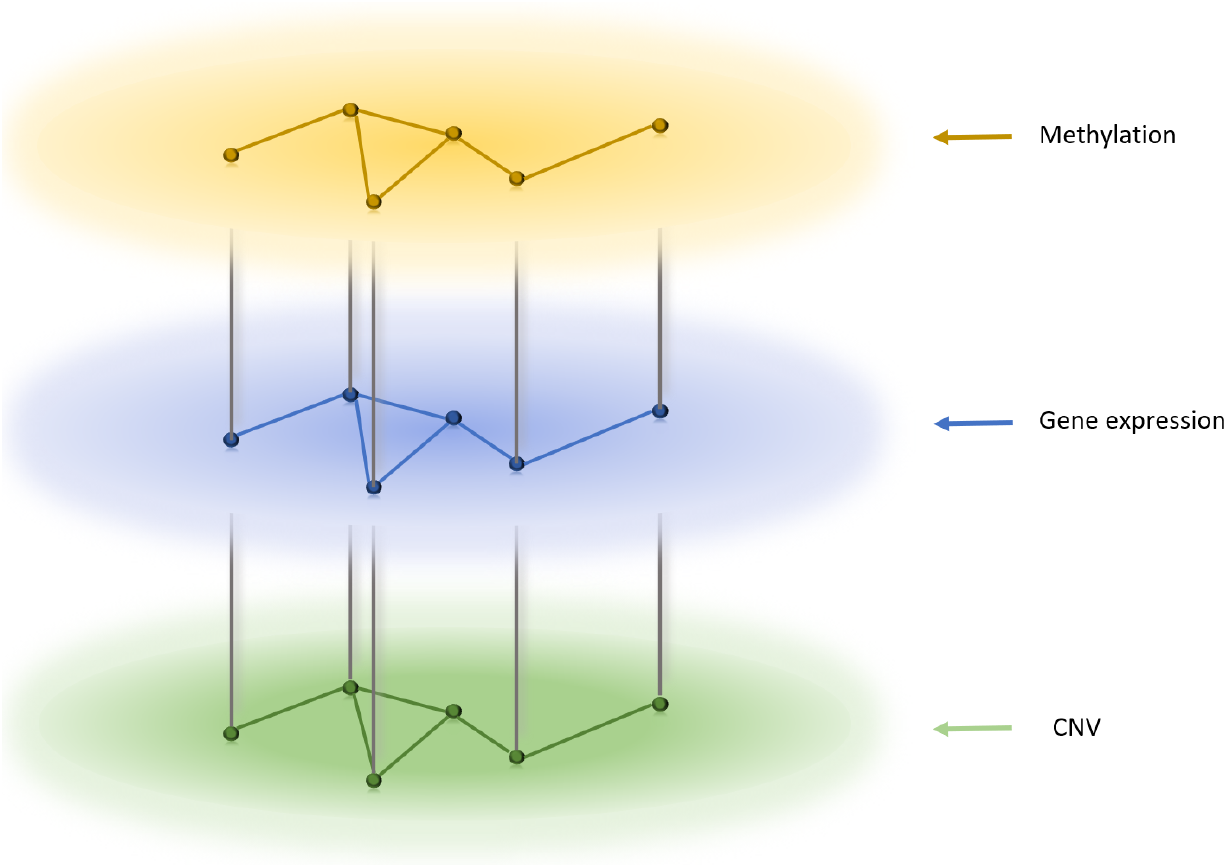
Graph structure for TCGA

**Figure 2:**
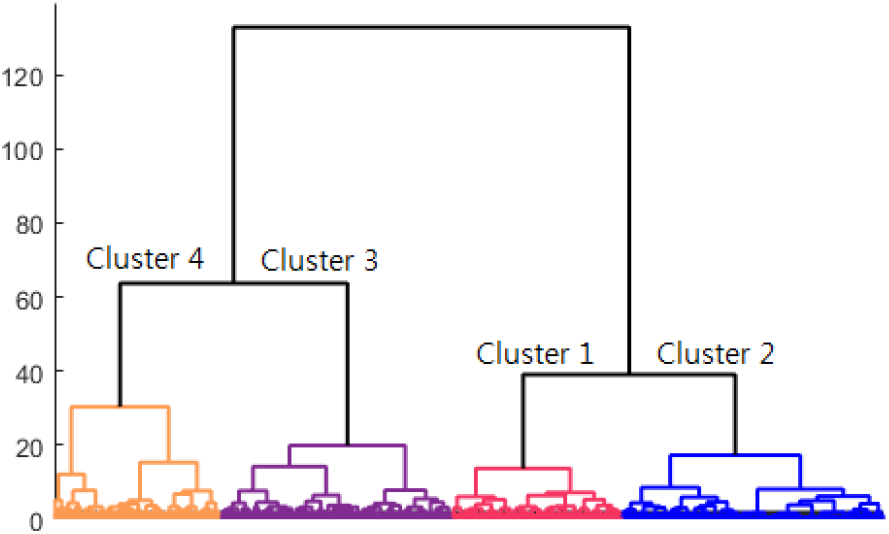
Clustering results employing vector-valued Wasserstein distance on METABRIC data.

Based on the dendrogram and the number of intrinsic molecular subtypes in breast cancer, 4 clusters were chosen for further analysis. Kaplan-Meier analysis with log-rank test (without NA sanples) resulted in a statistically significant survival difference among clusters with log-rank *p <* 0.0001(Figure 3).

**Figure 3:**
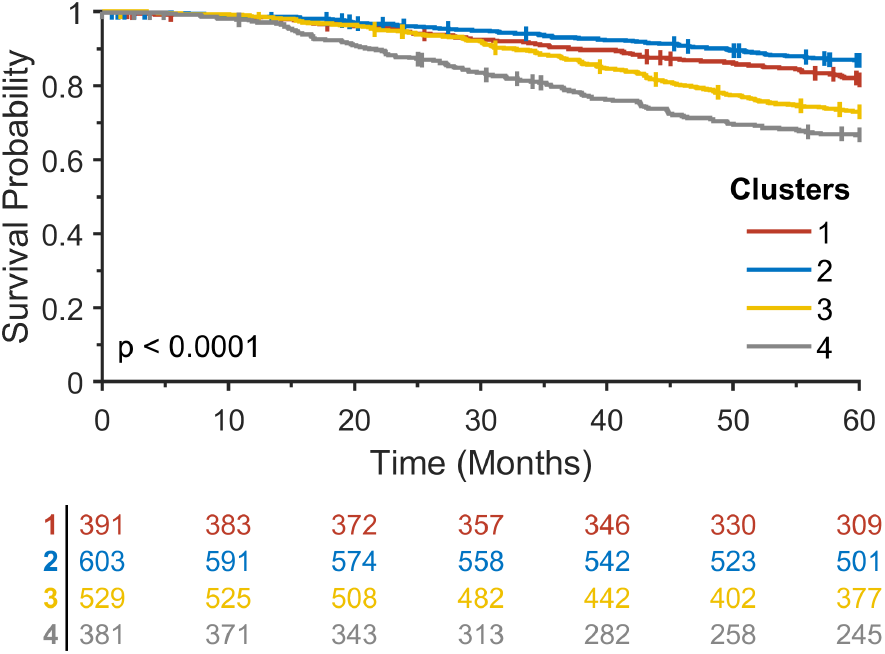
Kaplan-Meier analysis for four clusters that resulted from a hierarchical clustering method on the vector-valued Wasserstein distance matrix in the METABRIC study.

The clustering results were compared with PAM50 and Claudin-low subtypes[22], and the associations were assessed using a chi-squared test, resulting in p *<* 0.0001 as shown in Table 1. Clusters 1 and 2 were enriched for Luminal A subtype. Cluster 3 was enriched for luminal A and B subtypes, and cluster 4 was more enriched for basal subtype.

**Table 1:** Comparison between PAM50 along with Claudin-low subtypes and clusters identified by the proposed method. NA: not available.

| Cluster | Luminal A | Luminal B | Her2 | Basal | Claudin-low | Normal-like | NA |
| --- | --- | --- | --- | --- | --- | --- | --- |
| 1 | 189 | 94 | 33 | 16 | 24 | 32 | 3 |
| 2 | 316 | 78 | 32 | 18 | 95 | 62 | 2 |
| 3 | 147 | 205 | 85 | 31 | 27 | 33 | 1 |
| 4 | 27 | 84 | 70 | 134 | 53 | 13 | 0 |

**Table 2:** Comparison of 22 CIBERSORT immune cell types among four clusters in METABRIC, showing mean (standard deviation) values.

| Immune cell types | Cluster 1 | Cluster 2 | Cluster 3 | Cluster 4 | P-value |
| --- | --- | --- | --- | --- | --- |
| B cells naive | 0.008 (0.017) | 0.008 (0.018) | 0.008 (0.019) | 0.007 (0.017) | 0.8240 |
| B cells memory | 0.03 (0.036) | 0.035 (0.047) | 0.025 (0.031) | 0.026 (0.031) | **3.60E-05** |
| Plasma cells | 0.17 (0.098) | 0.169 (0.097) | 0.172 (0.093) | 0.165 (0.089) | 0.7405 |
| T cells CD8 | 0.045 (0.05) | 0.044 (0.048) | 0.041 (0.049) | 0.042 (0.047) | 0.5181 |
| T cells CD4 naive | 0.014 (0.034) | 0.017 (0.039) | 0.013 (0.028) | 0.009 (0.025) | **0.0028** |
| T cells CD4 memory resting | 0.055 (0.06) | 0.057 (0.058) | 0.052 (0.057) | 0.044 (0.053) | **0.0068** |
| T cells CD4 memory activated | 0.001 (0.005) | 0.001 (0.007) | 0.001 (0.005) | 0.001 (0.007) | 0.6263 |
| T cells follicular helper | 0.057 (0.034) | 0.053 (0.035) | 0.055 (0.035) | 0.068 (0.036) | **3.86E-10** |
| T cells regulatory (Tregs) | 0.016 (0.02) | 0.014 (0.02) | 0.017 (0.021) | 0.017 (0.021) | 0.0771 |
| T cells gamma delta | 0.057 (0.045) | 0.056 (0.044) | 0.059 (0.044) | 0.063 (0.044) | 0.1604 |
| NK cells resting | 0.003 (0.013) | 0.005 (0.016) | 0.003 (0.011) | 0.003 (0.013) | 0.1057 |
| NK cells activated | 0.029 (0.026) | 0.025 (0.026) | 0.029 (0.025) | 0.031 (0.028) | **0.0028** |
| Monocytes | 0.017 (0.024) | 0.021 (0.027) | 0.015 (0.022) | 0.019 (0.029) | **0.0018** |
| Macrophages M0 | 0.106 (0.095) | 0.104 (0.092) | 0.13 (0.098) | 0.158 (0.094) | **3.50E-19** |
| Macrophages M1 | 0.076 (0.042) | 0.069 (0.044) | 0.081 (0.043) | 0.103 (0.051) | **1.69E-28** |
| Macrophages M2 | 0.159 (0.089) | 0.162 (0.095) | 0.147 (0.077) | 0.126 (0.067) | **6.74E-11** |
| Dendritic cells resting | 0.003 (0.01) | 0.005 (0.013) | 0.003 (0.008) | 0.005 (0.014) | **0.0323** |
| Dendritic cells activated | 0.003 (0.01) | 0.004 (0.013) | 0.005 (0.017) | 0.009 (0.023) | **3.89E-07** |
| Mast cells resting | 0.147 (0.104) | 0.148 (0.11) | 0.144 (0.1) | 0.101 (0.082) | **4.26E-13** |
| Mast cells activated | 0.001 (0.004) | 0.001 (0.005) | 0.001 (0.004) | 0.001 (0.008) | 0.1828 |
| Eosinophils | 0 (0) | 0 (0.001) | 0 (0.001) | 0 (0) | 0.9508 |
| Neutrophils | 0.001 (0.005) | 0.001 (0.004) | 0.001 (0.006) | 0.001 (0.002) | 0.0684 |

For the four clusters identified, 22 CIBERSORT immune cell types were compared using one-way analysis of variance (ANOVA) test. Twelve immune cell types were statistically significantly different among the four clusters. The top two significant immune cell types were Macrophages M0 and Macrophages M1, for which Cluster 4 had the highest values with mean=0.158 (standard deviation [SD]=0.094) and 0.103 (0.051), respectively.

### 3.2 TCGA data analysis

To validate the proposed method, we further analyzed multi-omics data in the TCGA breast cancer study, including gene expression, CNV, and methylation data. The clustering results are shown in Figure 4. Similar to the METABRIC analysis, four clusters were chosen for further analysis. Kaplan-Meier analysis with log-rank test resulted in a statistically significant survival difference among clusters with log-rank p = 0.0088 (Figure 5).

**Figure 4:**
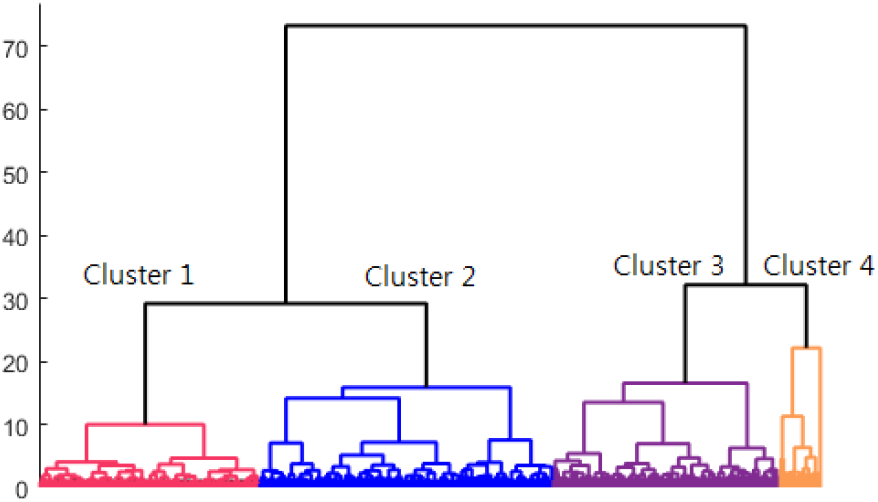
Clustering results employing vector-valued Wasserstein distance on TCGA data.

**Figure 5:**
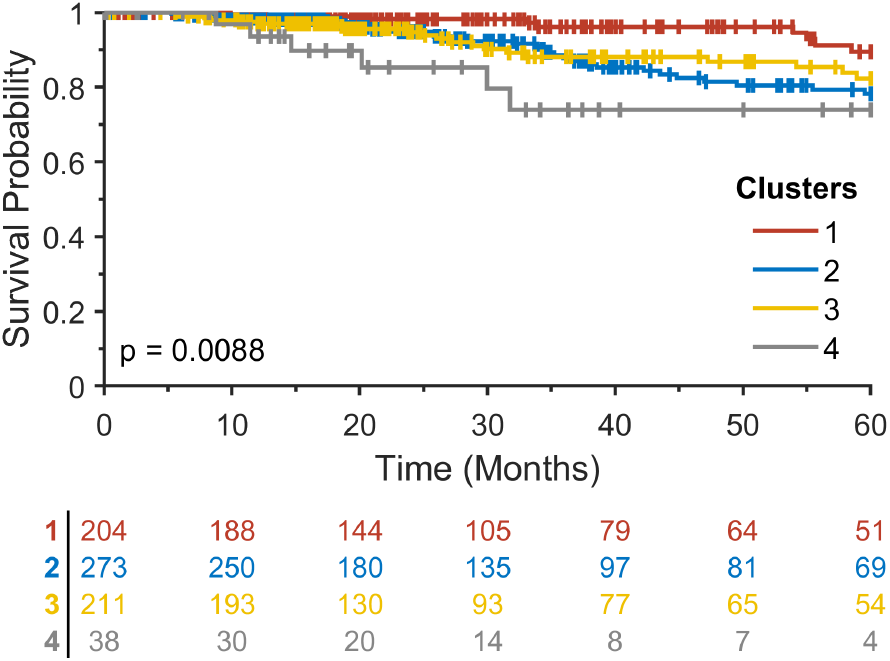
Kaplan-Meier analysis for four clusters that resulted from a hierarchical clustering method on the vector-valued Wasserstein distance matrix in the TCGA breast cancer study.

The clustering results were compared with PAM50. The associations were assessed using a chi-squared test, resulting in p *<* 0.0001 as shown in Table 3. Clusters 1 and 2 were enriched for luminal A and B subtypes. Cluster 4 was more enriched for basal subtype.

**Table 3:** Comparison between PAM50 and clusters identified by the proposed method. NA: not available.

| Cluster | Luminal A | Luminal B | Her2 | Basal | Normal-like | NA |
| --- | --- | --- | --- | --- | --- | --- |
| 1 | 184 | 9 | 0 | 0 | 11 | 0 |
| 2 | 131 | 46 | 22 | 56 | 15 | 3 |
| 3 | 66 | 73 | 19 | 45 | 4 | 4 |
| 4 | 2 | 7 | 2 | 25 | 1 | 1 |

For the four clusters identified, 22 CIBERSORT immune cell types were compared using the one-way ANOVA test. Fifteen immune cell types were statistically significantly different among the four clusters. The most significant immune cell type was Macrophages M0 with p =1.51E-08.

| Immune cell types | Cluster 1 | Cluster 2 | Cluster 3 | Cluster 4 | P-value |
| --- | --- | --- | --- | --- | --- |
| B cells naive | 0.068 (0.045) | 0.05 (0.046) | 0.045 (0.046) | 0.033 (0.037) | **2.29E-07** |
| B cells memory | 0.007 (0.017) | 0.015 (0.031) | 0.01 (0.021) | 0.017 (0.025) | **0.0005** |
| Plasma cells | 0.049 (0.052) | 0.037 (0.045) | 0.046 (0.052) | 0.032 (0.038) | **0.0341** |
| T cells CD8 | 0.106 (0.057) | 0.108 (0.064) | 0.093 (0.061) | 0.094 (0.068) | **0.0331** |
| T cells CD4 naive | 0 (0.004) | 0 (0.001) | 0.002 (0.011) | 0 (0.002) | 0.0662 |
| T cells CD4 memory resting | 0.135 (0.076) | 0.122 (0.074) | 0.098 (0.071) | 0.068 (0.069) | **1.23E-06** |
| T cells CD4 memory activated | 0 (0.002) | 0.004 (0.012) | 0.003 (0.011) | 0.003 (0.009) | **0.0014** |
| T cells follicular helper | 0.063 (0.039) | 0.073 (0.04) | 0.07 (0.045) | 0.093 (0.068) | **0.0003** |
| T cells regulatory (Tregs) | 0.014 (0.02) | 0.026 (0.03) | 0.02 (0.024) | 0.015 (0.02) | **8.15E-06** |
| T cells gamma delta | 0.003 (0.01) | 0.003 (0.01) | 0.002 (0.007) | 0.004 (0.012) | 0.4291 |
| NK cells resting | 0.003 (0.01) | 0.006 (0.013) | 0.005 (0.011) | 0.012 (0.019) | **7.92E-05** |
| NK cells activated | 0.02 (0.024) | 0.019 (0.023) | 0.021 (0.025) | 0.017 (0.023) | 0.8629 |
| Monocytes | 0.02 (0.023) | 0.016 (0.019) | 0.015 (0.026) | 0.016 (0.018) | **0.0314** |
| Macrophages M0 | 0.056 (0.1) | 0.087 (0.105) | 0.116 (0.132) | 0.177 (0.165) | **1.51E-08** |
| Macrophages M1 | 0.055 (0.032) | 0.069 (0.045) | 0.057 (0.04) | 0.058 (0.054) | **0.0014** |
| Macrophages M2 | 0.271 (0.119) | 0.273 (0.128) | 0.301 (0.121) | 0.286 (0.13) | 0.1177 |
| Dendritic cells resting | 0.021 (0.031) | 0.015 (0.025) | 0.011 (0.024) | 0.005 (0.012) | **0.0003** |
| Dendritic cells activated | 0.002 (0.007) | 0.004 (0.013) | 0.009 (0.026) | 0.021 (0.06) | **3.58E-07** |
| Mast cells resting | 0.096 (0.071) | 0.063 (0.062) | 0.064 (0.075) | 0.042 (0.044) | **1.41E-07** |
| Mast cells activated | 0.007 (0.024) | 0.005 (0.02) | 0.011 (0.036) | 0.004 (0.009) | 0.4921 |
| Eosinophils | 0.001 (0.003) | 0 (0.003) | 0 (0.002) | 0 (0) | 0.8402 |
| Neutrophils | 0.003 (0.008) | 0.002 (0.005) | 0.004 (0.007) | 0.003 (0.008) | 0.1433 |

Nine immune cell types were significantly different in both METABRIC and TCGA studies: B cells memory, T cells CD4 memory resting, T cells follicular helper, Monocytes, Macrophages M0, Macrophages M1, Dendritic cells resting, Dendritic cells activated, and Mast cells resting (Figure 6). Three immune cell types, including T cells CD4 naive, NK cells activated, and Macrophages M2, showed statistical significance in METABRIC alone, whereas six immune cell types, including B cells naive, Plasma cells, T cells CD8, T cells CD4 memory activated, T cells regulatory (Tregs), and NK cells resting, showed statistical significance in TCGA alone.

**Figure 6:**
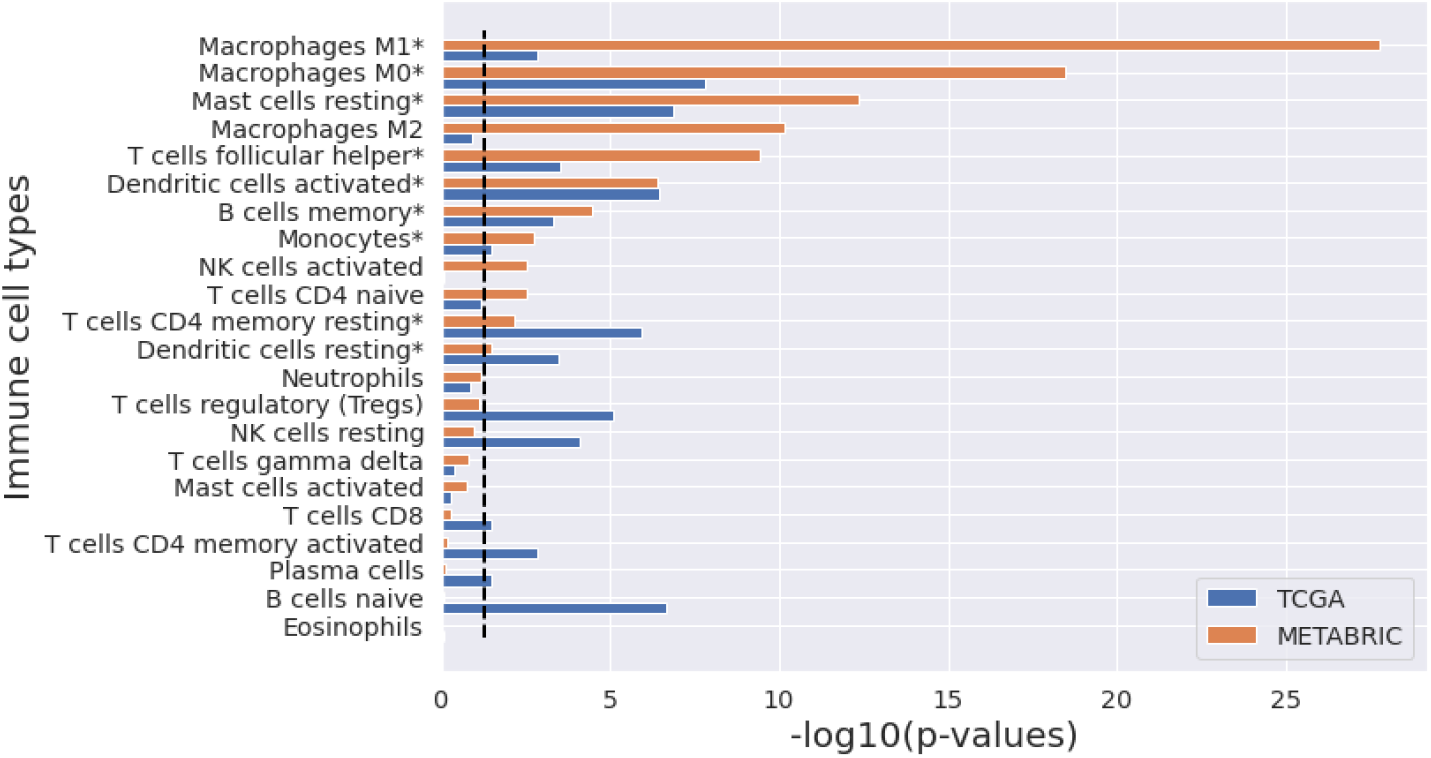
Comparison of 22 immune cell types in CIBERSORT among four clusters identified.

This indicates that the immune cell types are statistically significant (ANOVA p *<* 0.05) in both METABRIC and TCGA studies. The black dot line indicates -log10(0.05).

## 4 Discussion

The treatment of multi-omic biological data in a vector-valued manner may provide new insights for understanding the biological mechanisms of cancer biology, using complementary information offered by individual omics types. vWCluster is a data analysis methodology based on OMT theory, which enables the integration of multi-omics data in a vector-valued form, represented by multiple layers in a network. The integrated invariant measure was further employed to identify cancer subtypes. We applied this method to the two largest breast cancer studies, METABRIC and TCGA. Clusters identified showed significantly different survival rates in both studies.

CIBERSORT scores, consisting of 22 immune cell types, were further compared among the identified clusters. CIBERSORT employs gene expression profiles from a set of 547 genes to predict 22 immune cell types, using support vector regression [21]. ANOVA tests revealed that 9 immune cell types were commonly statistically significant in both studies, indicating that the tumor immune microenvironment varies in the identified clusters and this is associated with the difference in survival in breast cancer patients.

Kaplan-Meier analysis was performed for intrinsic molecular subtypes in the METABRIC and TCGA breast cancer studies (Figure 7). As in Kaplan-Meier analysis for the clusters identified by our method in METABRIC, an extremely significant survival difference was found among intrinsic subtypes with a log-rank p-value *<* 0.0001. By contrast, for the TCGA breast cancer cohort, our method resulted in much better statistical significance with a log-rank p=0.0088 compared to marginal statistical significance with a log-rank p=0.0291 among intrinsic subtypes in TCGA, suggesting the potential of our proposed method to identify new subtypes in cancer and further stratify patients at high risk of mortality.

**Figure 7:**
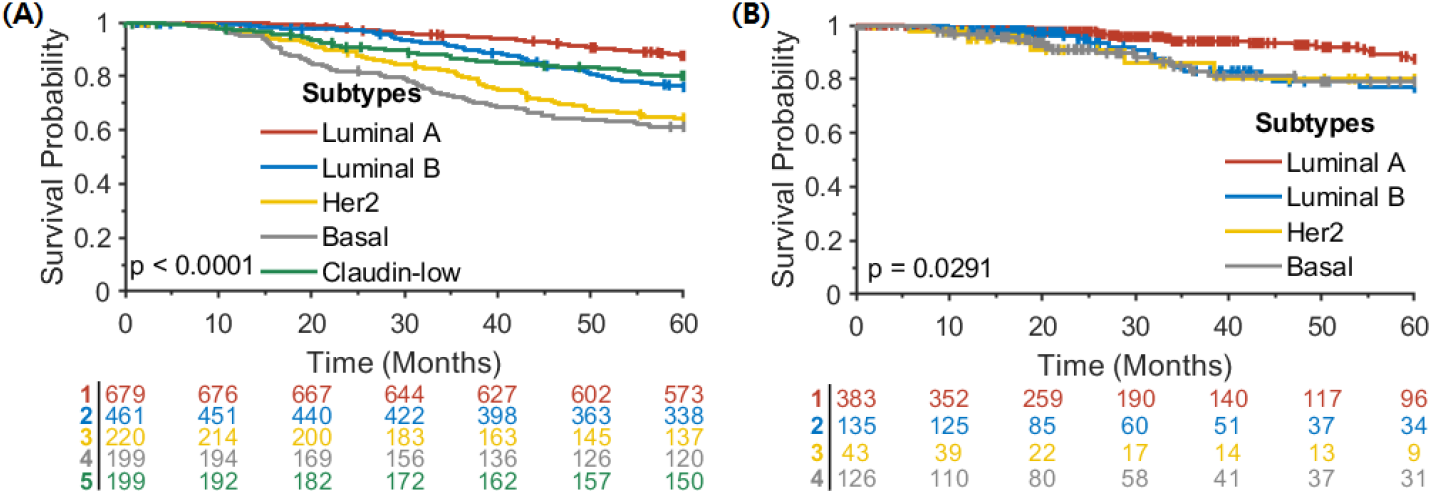
Kaplan-Meier analysis for intrinsic molecular subtypes in (A) METABRIC and (B) TCGA with no normal-like samples.

## 5 Conclusion

We proposed a multi-omics data integration and clustering method, called vWCluster, based on the vector-valued Wasserstein distance. In this method, individual omics types represented as multiple layers in a network can be efficiently integrated, considering the biological interactions of biomarkers and providing complementary biological information. The formulation of vWCluster treats the data vectorially, which potentially minimizes information loss. vWCluster is flexible and applicable to the integration of multi-modal data including imaging and genomic data, which is a research direction we plan to explore in the future.

## 6 Acknowledgments

This study was supported by AFOSR grants (FA9550-17-1-0435, FA9550-20-1-0029), NIH grants (R01-AG048769, R21-CA234752), NIH/NCI Cancer Center Support grant (P30 CA008748), and a grant from Breast Cancer Research Foundation (grant BCRF-17-193).

